# FONDUE: Robust resolution-invariant denoising of MR Images using Nested UNets

**DOI:** 10.1101/2023.06.04.543602

**Authors:** Walter Adame-Gonzalez, Aliza Brzezinski-Rittner, M. Mallar Chakravarty, Reza Farivar, Mahsa Dadar

## Abstract

Recent human neuroimaging studies tend to have increased magnetic resonance image (MRI) acquisition resolutions, seeking finer levels of detail and more accurate brain morphometry. However, higher-resolution images inherently contain greater amounts of noise contamination, leading to poorer quality brain morphometry if not addressed adequately. This study proposes a novel, robust, resolution-invariant deep learning method to denoise structural human brain MRIs. We explore denoising of T1-weighted (T1w) brain images from varying field strengths (1.5T to 7T), voxel sizes (1.2mm to 250µm), scanner vendors (Siemens, GE, and Phillips), and diseased and healthy participants from a wide age range (young adults to aging individuals). Our proposed Fast-Optimized Network for Denoising through residual Unified Ensembles (FONDUE) method demonstrated stable denoising capabilities across multiple resolutions with performance comparable to the state-of-the-art methods. FONDUE was capable of denoising 0.5mm^3^ isotropic T1w images in under 3 minutes on an NVIDIA RTX 3090 GPU using less than 8GB of video memory. We have also made the repository of FONDUE as well as its trained weights publicly available on: https://github.com/waadgo/FONDUE.

## 1. Introduction

Magnetic Resonance Imaging (MRI) is a non-invasive imaging modality that has widespread applications in clinical diagnosis and research. It offers distinct anatomical information compared to other imaging modalities such as Computed Tomography (CT) or ultrasound, as it can be adjusted to achieve high contrast between various tissue types (e.g., in brain tissue between grey matter, GM, and white matter, WM) and different pathologies (e.g. tumors and lesions).

MR images are affected by noise that originates from various sources such as stochastic variation, physiological processes, eddy currents, artifacts from the differing magnetic field susceptibilities of adjacent tissues, rigid body motion, nonrigid motion, etc. [1]. There are several methods for quantifying relative levels of noise in MRI, with signal-to-noise ratio (SNR) being one of the most commonly used metric of image quality, measuring the ratio of true signal to noise. SNR in MRI is usually obtained by computing the difference between a region of interest (ROI) selected from an image area containing anatomical information and the background, which should consist of only air, and thus, any signal present in it should be noise. The computed difference is divided by the standard deviation of the signal present in the background.

High-resolution neuroimaging can provide more anatomical details with higher accuracy. This is particularly important when studying relatively small brain regions that might not be easily detected in lower-resolution MRIs. Recent large-cohort studies have started to explore sub-millimeter voxel sizes, such as the Human Connectome Project (HCP) [2], in which 1200 healthy young adults were scanned at sub-millimeter resolutions. However, decreasing the voxel size to obtain high-resolution MRIs leads to lower SNR [3]. Therefore, the usage of high-resolution MRIs for brain morphometry and image segmentation should be carefully planned since otherwise, the results will inevitably contain considerable amounts of noise contamination, hindering the quality of MRI segmentations as the true anatomy signal on the image gets corrupted [4].

To enhance SNR in MRI, one can: use spin-echo sequences instead of gradient echo, use a smaller bandwidth, use surface coils and increase the number of excitations, increase signal by decreasing the echo time (TE), or increase the repetition time (TR) [3]. Scan parameters can also be modified to increase SNR (assuming all other factors remain constant) by increasing the field of view (FOV), reducing matrix size, and increasing the slice thickness. One may also increase SNR by averaging multiple acquisitions of the same sequence. However, this comes at the cost of a significant increase in acquisition time and requires the participant to spend large amounts of time inside the scanner, something that might not be feasible in clinical settings. Finally, averaging multiple acquisitions requires co-registration of the images since participants are more likely to move inside the scanner as scanning time increases.

Unlike the above-mentioned techniques which are mainly related to adjusting the acquisition protocol and entail an increase in the time and cost associated with acquisition, denoising methods can potentially generate high-SNR high-resolution images without adding to acquisition time and costs. Therefore, denoising methods are a promising way to address low-SNR in high-resolution MRIs.

Most denoising tools that are currently used in the neuroimaging field involve non-deep learning (NDL) methods that are based on different techniques such as sparseness of voxel patches (e.g. Adaptive Optimized Non-Local Means; AONLM), averaging nonlocal patches (e.g. Non-Local Means; NLM), and Principal Components Analysis to extract the true-signal information using a pre-filtered image as a guide (Pre-filtered Rotationally Invariant Non-Local Principal Component Analysis; PRI-NLPCA) [5]–[13]. NLM is a filtering method that averages similar pixels located in different regions of the image. It is based on the idea of finding similar pixels in the whole image based on their neighborhood. Thus, a given pixel is averaged with similar pixels using a similarity parameter based on an exponential function that includes a Gaussian-weighted Euclidean distance [5]. Block-Matching 4D denoising (BM4D) [6] is a filtering method in which a group of 3D sub-units (blocks) extracted from a 3D image are grouped based on a hard-threshold similarity metric and then stacked together in 4D units. Analogously to NLM, BM4D uses non-local blocks during the matching stage followed by a collaborative filtering stage in which through a “joint four-dimensional transform” they compute similarity within each block and within each 4D stack of blocks, Wiener filtering, and applying the inverse joint four-dimensional transform after aggregation of previously computed adaptive weights.

AONLM is an NLM-based denoising algorithm that can deal with spatially varying noise typically present in parallel imaging such as the one present in generalized auto-calibrating partially parallel acquisitions (GRAPPA). Furthermore, AONLM uses a block-wise denoising approach instead of voxel-wise denoising used by NLM, significantly improving the processing time with a negligible decrease in filtering quality. Finally, they proposed an adaptation that allows for filtering of Rician-distributed non-stationary noise which uses a local smoothing factor (sigma) rather than a global one [7].

PRI-NLPCA [8] is a modified and improved version of Pre-filtered Rotationally Invariant Non-Local Means (PRI-NLM) [9]. Both algorithms use a modification of the standard NLM algorithm to compute the denoised image by introducing a Rotationally Invariant similarity metric to compute the non-local means. Therefore, instead of using Euclidean distance to compute similarity, similar patches can be used for computing means whether their orientation is similar or not. However, instead of using the noisy image to compute the similarity within the search window, a pre-filtered image is used as the guide image to compute pixel similarities. This helps the computation of the similarity between pixels in the rotationally invariant approach since it is known that rotationally invariant descriptors are sensitive to noise.

Due to their nature, NDL denoising methods do not require target images, nor extensive model training to create a functioning method. However, most NDL denoising methods suffer from a trade-off between resolution/denoising accuracy and processing time. For instance, in PRI-NLPCA [8] they explored the optimal value for the sliding window overlap while exploring stride (the number of voxels that the sliding window moves from iteration to iteration) values from 1 to 3 (maximum to minimum overlap). As expected, the best performance was obtained when the overlap was maximum while the processing time was the longest since more computations were performed as more information was used. Hence, the potential widespread implementation in a clinical environment or in large-cohort research studies of NDL denoising methods that use this type of sliding window approach might be limited especially when high-resolution MRIs are being used.

Deep Learning (DL) denoising methods are techniques that use Machine Learning (ML) principles to generate a function capable of mapping a representation to a certain given image. Most DL methods employ Convolutional Neural Networks (CNNs), which require training datasets with pairs of images: noisy images and their corresponding noise-free images. These DL denoising methods aim to learn the transformations from the noisy image to the noise-free image. This is achieved by exploiting the capabilities of CNNs as universal function approximators through backpropagation. It is important to note that these approaches have recently benefited from the use of dedicated graphical processing units (GPUs), which make them several orders of magnitude faster than NDL algorithms, even when the DL methods are used on the same CPU as the ones used by NDL instead of a dedicated GPU. Moreover, DL methods can outperform NDL methods in many medical image-denoising tasks [10]–[14]. Despite the aforementioned advantages of using DL approaches for image denoising, they face the following drawbacks:

- Requiring large amounts of curated data: Having sufficient and representative training and validation data is crucial for optimal training. In other words, having adequate variability within the training data enables the network to generalize to unseen data instead of overfitting (i.e., performing well only for a small subset of data similar to its training dataset and poorly when applied to new previously unseen data). In general, the more representative the data used for training is, the better the performance that can be achieved.
- DL denoising methods tend to introduce different levels of undesired blurring on high-frequency details (e.g. cerebellar GM and WM-GM interfaces) due to their convolutional nature and the use of operations such as pooling. This can potentially lead to undesired loss of anatomical information or changes in the anatomy [15].
- Need to train on fixed-resolution images: In order to optimally train the weights in a CNN, images with different resolutions must be re-scaled to match the requirements of the network so that the anatomical structures are all within approximately the same scale in pixel dimensions. Therefore, if it is necessary to denoise images at several resolutions, different networks should be trained for the different resolutions that one would like to denoise images at. This limitation can significantly restrict the applicability of DL denoising methods in practice [16], [17].

Different network architectures have been proposed in the literature for image denoising. From the popular DnCNN [14] and its adaptation to MRI denoising MCDnCNN [18] which are architectures that resemble a ResNet [19], to encoder-decoder-like architectures similar to the original U-Net [20] which was proposed for segmentation tasks, adapted to perform volumetric medical image denoising, like in RDUNet [11]. It has been shown, however, that a densely connected and nested version of U-Net, i.e., U-Net++ [21] can outperform the original U-Net version in segmentation tasks for medical images. We may then think that this type of architecture can better learn relevant image features that are important for segmentation tasks. Such an improvement in learning at the latent space level can also be used for other tasks such as regression, like image denoising. Finally, residual CNNs and GANs have shown to perform better in denoising tasks than non-residual approaches, since it is easier to learn the latent noise of the noisy input than learning to infer the clean image from it [14].

In this study, we propose a new, robust method for denoising structural brain MRIs by employing a DL approach that can be used to denoise any images of any resolution while maintaining the structural integrity of the high-frequency response. Furthermore, our proposed method will allow users to obtain denoised images with similar peak SNR (PSNR) values to state-of-the-art denoising methods with faster processing times.

## 2. Methods

### 2.1. Datasets

In order to obtain a robustly trained and generalizable CNN, we need to use brain images with enough feature variability for both training and validation. Therefore, for the methodology design, we included datasets with different magnetic field strengths (1.5T and 3T), scanner vendors (SIEMENS, Phillips, and GE), acquisition protocols, resolutions (from over a millimetre voxel sizes to sub-millimetre voxel sizes), subject ages (from young adults to aged individuals), and diseases (healthy, Mild Cognitive Impairment, Alzheimer’s Disease, Parkinson’s Disease, Schizophrenia, etc.). We also included a balanced number of female and male participants in our dataset. This careful pre-selection of the training set can have a major impact in terms of generalization capabilities of our trained network. The following section provides a brief description of the datasets that were used in the training of our proposed model. Table 1 includes a summary of the characteristics of the images included for each dataset.

**Table 1.** Summary of the datasets used to train and validate FONDUE.

| Study | Cohort(s) | Field Strength | Scanner Model | Repetition time [ms] | Echo time [ms] | Flip Angle [°] | Voxel dimension [mm] | Training sample [N] | Validation sample [N] | Generalization sample [N] |
| --- | --- | --- | --- | --- | --- | --- | --- | --- | --- | --- |
| ABIDE-II | AS | 3T | Philips, GE, Siemens | 3 | 3.9 | 8 | Multiple | 169 | 14 | 20 |
| ADNI1 | HA, MCI, AD | 1.5T, 3T | Philips, GE, Siemens | ~7-8 | ~2-4 | ~8-12 | 1.0×1.0×1.2 | 131 | 11 | 20 |
| HCP | HA | 3T | Siemens | 2.4 | 2.14 | 8 | 0.7×0.7×0.7 | 174 | 15 | 20 |
| IXI | HA | 1.5T | GE, Philips | 9.6 | 4.6 | 8 | 1.0×1.0×1.0 | 407 | 25 | 20 |
| LA5c | HA, SZ, ADHD | 3T | Siemens | 1.9 | 2.26 | 7 | 1.0×1.0×1.0 | 89 | 16 | 20 |
| MIRIAD | AD | 1.5T | GE | 15 | 5.4 | 15 | 0.937×0.937×1.5 | 48 | 7 | 14 |
| Custom_0.5 | HA | 3T | Siemens | 2750 | 2.2, 4.63 | N/A | 0.5×0.5×0.5 | 12 | 3 | 3 |
| UH_250 | HA | 7T | Siemens | 3580 | 2.41 | 5 | 0.25×0.25×0.25 | 0 | 0 | 1 |
| UH_500 | HA | 7T | Siemens | 2740 | 3.24 | 5 | 0.5×0.5×0.5 | 0 | 0 | 1 |
| UH_1000 | HA | 7T | Siemens | 2500 | 1.91 | 5 | 1.0×1.0×1.0 | 0 | 0 | 1 |

Acronyms: AS = Autism Spectrum, HA = Healthy Adults, MCI = Mild Cognitive Impairment, AD = Alzheimer's Disease, ADHD = Attention-Deficit / Hyperactivity Disorder, SZ = Schizophrenia

#### ABIDE-II

The Autism Brain Imaging Data Exchange II [22] is a dataset including MRIs from individuals with autism spectrum disorder as well as matched controls. Study sites used either 3D magnetization prepared rapid gradient echo (MPRAGE) sequences or vendor-specific custom variants. While 3T images from GE, Philips, and Siemens vendors were available in this dataset, images acquired with a Philips Achieva scanner will be used for this project in order to balance the number of samples from each vendor and to have native images of varying resolutions, mostly non-isotropic, varying from over to sub-millimetre.

#### ADNI-1

The Alzheimer’s Disease Neuroimaging Initiative 1 [23] includes data from normal aging individuals as well as people with subjective cognitive decline, mild cognitive impairment, and Alzheimer’s dementia such as structural and functional MRI, positron-emission tomography (PET), as well as biological biomarkers and clinical assessments; it was collected across 59 study sites and includes 1.5T and 3T MRIs from different vendors. From the ADNI-1, we included MRI scans from GE, Siemens, and Philips scanners with voxel resolution of 1.0×1.0×1.2 mm^3^. Due to the numerous testing sites that acquired MRIs for ADNI-1, the acquisition parameters vary from site to site, providing us with a variety of parameter ranges presented in Table 1.

#### HCP

The Human Connectome Project Young Adult study [2], is a dataset comprised of brain images from 1200 healthy subjects ranging from 22 to 35 years old acquired using a 3T Siemens scanner with a sub-millimetre voxel size (isotropic 0.7 mm^3^).

#### IX

The IXI Dataset [24] includes MRI scans from 600 normal healthy subjects from three different hospitals in London, UK. Phillips 3T and 1.5T systems were used at Hammersmith and Guy’s Hospitals, respectively.

#### LA5c

The UCLA Consortium for Neuropsychiatric Phenomics LA5c Study [25] includes scans from 130 healthy controls, 43 individuals with diagnoses of adult Attention-Deficit Hyperactivity Disorder (ADHD), 49 individuals with bipolar disorder, and 50 individuals with schizophrenia. Neuroimaging data were acquired on a Siemens 3T scanner using an MPRAGE sequence.

#### MIRIAD

The Minimal Interval Resonance Imaging in Alzheimer’s Disease [26] is a longitudinal study containing T1 MRI scans of 46 patients with Alzheimer’s disease at intervals of 2, 6, 14, 26, 38 and 52 weeks, 18 and 24 months from baseline. The images were acquired with a GE 1.5T scanner using an inversion recovery prepared fast spoiled gradient recall sequence.

#### Custom_0.5

A custom dataset of 21 individuals ranging from 25 to 50 years old (11 females) acquired at the Montreal General Hospital (McGill University Research Center) [27] was also used as part of the training and validation corpus. We used images from 16 out of the 21 individuals that passed quality control. The dataset was acquired using a 3T Siemens Prisma scanner with a Multi-Echo MPRAGE (MEMPRAGE) sequence using a 64-channel head coil and online GRAPPA acceleration. Importantly, 14 out of the 16 individuals in CUSTOM_0.5 were scanned 6 times, allowing us to generate higher-quality average ground truth images. These individuals formed the dataset named CUSTOM_0.5_6rep, which was further divided for training and testing during training. A subset of 2 individuals (which was left out of the training set) from CUSTOM_0.5_6rep was further scanned another 14 times, generating a total of 20 scans for each of these 2 individuals, referred to as the CUSTOM_0.5_20rep dataset. CUSTOM_0.5_20rep subjects were scanned 20 times in 3 sessions separated by a few months. 6 scans were acquired on the first session and 7 scans on the next two sessions. The objective was to obtain the cleanest possible MRIs for each individual after averaging their respective 20 available scans to be used as gold standard for validating FONDUE.

#### UH_NNN

A novel dataset by [28] known for being the highest-resolution in-vivo structural MRI dataset available to date. UH-NNN consists of a single-subject dataset acquired with a 7T Siemens scanner at 0.25 mm^3^ isotropic voxel size. They collected 8 acquisitions at this resolution, using motion tracking correction, linear and non-linear co-registration using ANTs, non-homogeneity correction and slight denoising. They provide images of the same subject at three different resolutions: 0.25 mm^3^ isotropic (UH_250), 0.5 mm^3^ isotropic (UH_500) and 1.0 mm^3^ isotropic (UH_1000), with the acquisition parameters summarized in Table 1. We did not use any of these images for training, and performed qualitative validation of the performance of FONDUE in denoising a single repetition of the raw 7T images at the three different resolutions. We also quantitatively compared the performance of FONDUE against a commonly used NDL method in denoising a single repetition of UH_250. To generate the ground truth for these comparisons, we co-registered the 8 repetitions (rigid followed by nonlinear registration as was done in the original article, rescaled to 0.3 mm^3^ isotropic voxel size).

### 2.2. System

To build FONDUE, we used Python 3.9.7 along with PyTorch 1.10.0, *numpy* 1.22.2, and *nibabel* 3.2.1 to handle the usage of NIFTI and MGH imaging formats. The script was implemented using Anaconda’s Spyder editor in a Windows 11 system. Our hardware included an Intel Core i7-10700 as CPU, an NVIDIA RTX 3090 for GPU (24GB of VRAM), and 128GB of RAM memory.

### 2.3. Network Architecture

Current hardware memory limitations necessitate adopting either a 2.5D (a stack of 2D slices) approach or small 3D blocks (extracting regularly sized 3D blocks from the MRI volume) approach as it is currently impossible to process sufficiently large numbers of whole MRI volumes at once with CNNs. Therefore, the volumes must be divided into smaller pieces to fit in the computer’s memory. Several groups have previously successfully employed 2.5D approaches (instead of using 3D blocks) for both regression and segmentation tasks in MRI analysis. In [18], they used a stack of 5 slices of brain MRIs as input of their network called MCDnCNNg to remove noise from the middle slice. In FastSurferCNN [17] and FastSurferVINN [16], they used a stack of 7 slices from brain MRIs to perform semantic segmentation of the middle slice. These works showed that pilling up 5-7 slices can provide the network with enough spatial information (context) of the middle slice to increase the network performance while keeping memory usage as low as possible. Similar to these previous works, we adopted a 2.5D approach with a stack of 7 slices.

Figure 1 presents the architecture of the proposed DL denoising model. A stack of 7 consecutive slices (upper-left yellow box) are input into the network to denoise the middle slice (red contour). The stack is a tensor of size BS×C×H×W, where BS denotes the Batch Size, C are the channels (7 in this case), and H and W are the Height and Width of every image being input into the network. The blocks denoted by “i,j” in the figure refer to the convolutional blocks CBU_i,j_ (Figure 2). The blue-dotted arrows refer to the skip connections. The green dot denotes the maxout operation. The purple arrows denote a simple parsing to the maxout operation. The black arrows denote the bilinear interpolation operations. The orange arrows are the MaxPool2d operations plus indices parsing, and the golden arrows pointing upwards denote the MaxUnpool2d operations that use the indices previously parsed. Finally, the red arrows denote the final convolution step with a kernel of size BS×C×H×W and the yellow YN units with N={1,2,3,4,5,6} are the six noise candidates produced by all the sub-networks of FONDUE. The average of all YN images is subtracted from the noisy middle-slice to produce the denoised image.

**Figure 1:**
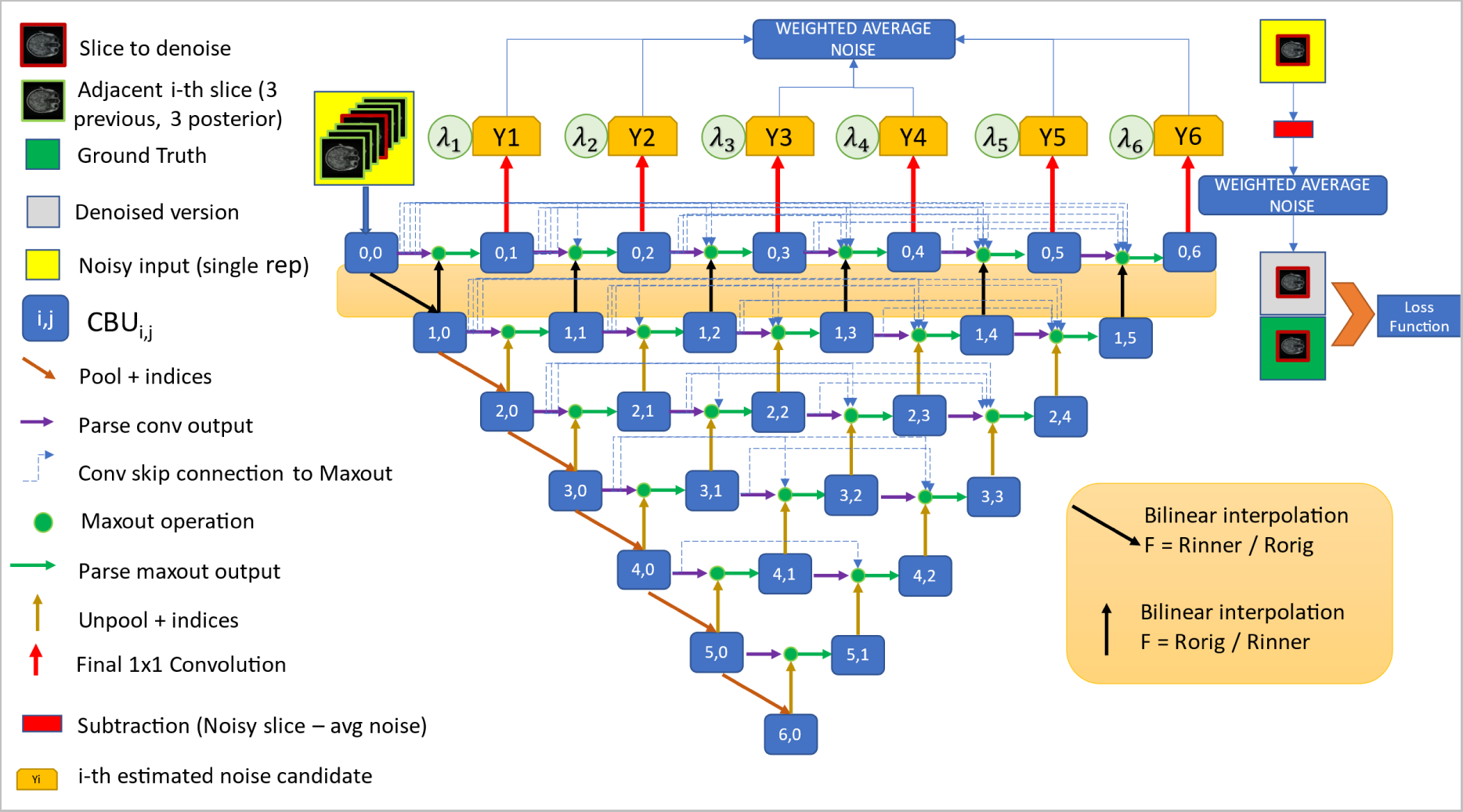
Network architecture of the proposed FONDUE method.

**Figure 2:**
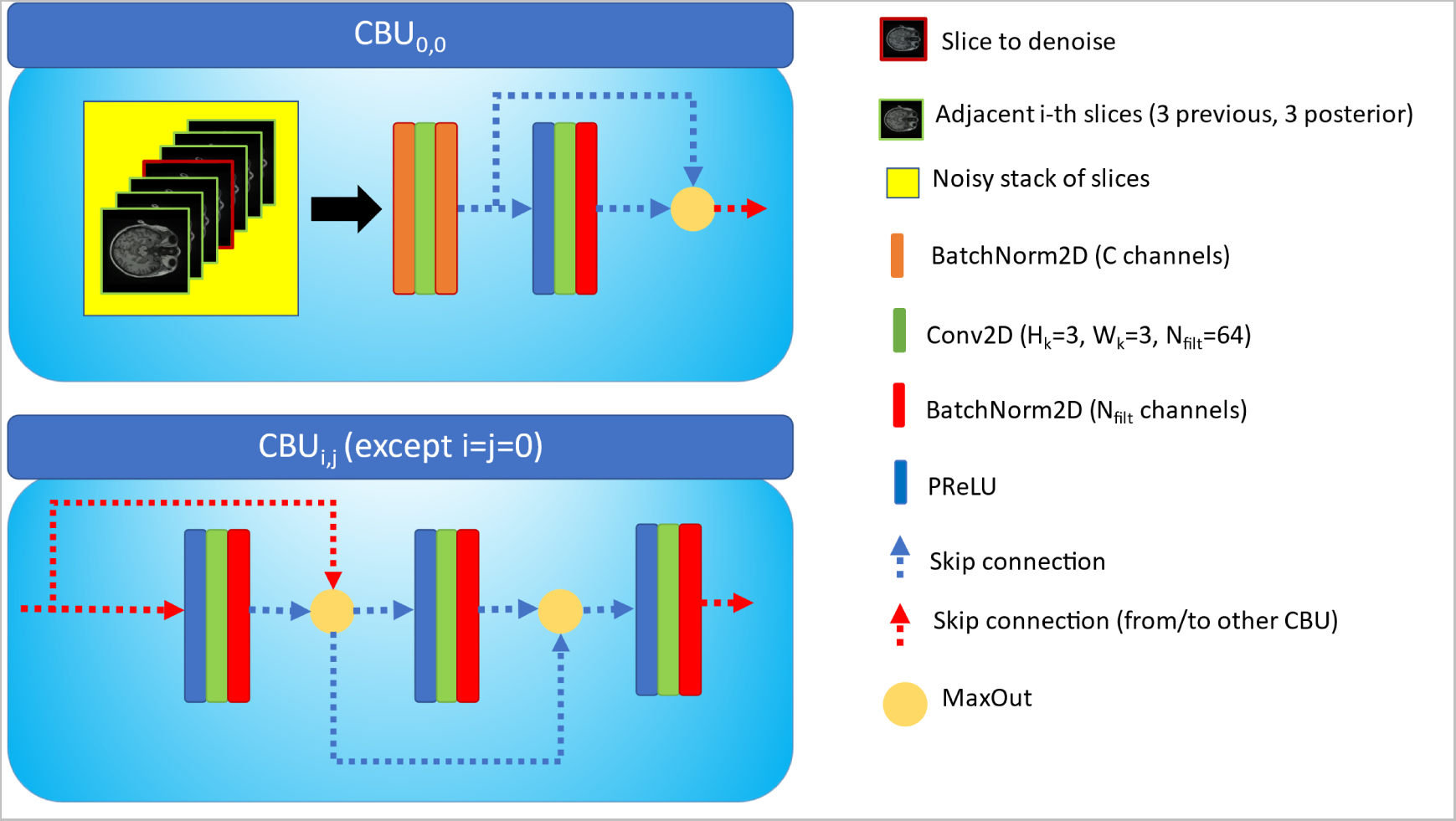
Convolutional Block Units design.

Similar to UNet++, we have adopted a densely connected U-shaped encoder-decoder architecture. As opposed to UNet and UNet++, we do not increase the number of filters as we go deeper into the network. We instead maintained the same number of filters at any position of the network, using *maxout* operations as proposed in [29] since it helps to significantly reduce the computational load while keeping good performance levels. We took advantage of *maxout*’s competitive nature to obtain a fast, reliable method that can be widely applicable and used. Additionally, we used convolutional block units (CBUs, Figure 2) similar to the Competitive Dense Blocks (CDB) proposed by [30].

After CBU0,0, we resample the feature maps by performing bi-linear interpolation using a factor F=Ro/Ri, where Ro is the original resolution in terms of the voxel size in millimeters, and Ri is the internal resolution of the network, which in this case was set to 1.0, but can be adjustable to any desired value. After filter bilinear interpolation, we feed the resulting feature maps into the CBU_1,0_. We then consecutively compute CBU_i,j_ with i={2, 3, 4, 5, 6} and j={0}, performing a MaxPooling operation with kernel size=(h, w) with h=2 and w=2, stride = 2 and padding = 0. Each MaxPooling operation reduces the dimensions of the filters to half. CBU_i,j_ with i={1, 2, 3, 4, 5} and j=1, 6-i is computed by performing the maxout operation between the MaxUnpool2d operation of CBU_i+1,j-1_ and all CBU_i,[0,j-1]_. Finally, we compute CBU_i,j_ with i=0 and j=[1, 6] by performing *maxout* of the bilinearly interpolated CBU_i+1,j-1_ (using a factor of 1/F, i.e., Ri/Ro) and all CBU_i,[0,j-1]_.

### 2.4. Weighted average

We trained 6 scalar parameters as weights for each of the noise candidates YN called *λN* with N ∈ {1, 2, 3, 4, 5, 6}. The final noise candidate Y is the weighted average of all 6 noise candidates Y_N_:

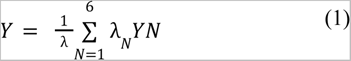

With:

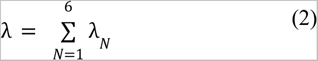

### 2.5. View aggregation

We trained FONDUE using only axial slices, but to obtain the denoised output for a new given image, we denoise using the three anatomical planes and average the results for the three denoised volumes. This process is known to help reduce block artifacts and visually improve image quality [16].

### 2.6. Loss function and image similarity metrics

Most denoising methods in the field have explored using Mean Absolute Error (MAE), also called L1 [11], [13], [31], and Mean Squared Error (MSE), also called L2 as loss functions [13], [18], [32].

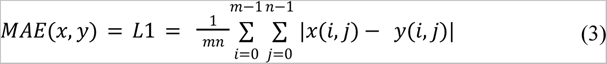

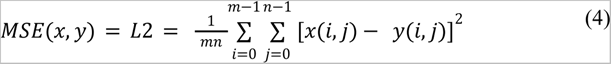

However, L1 and especially L2 loss functions can produce over smoothing (OS) in the output images when used alone [31]. To address OS, some authors have combined traditional L1 or L2 functions with structural similarity metrics such as Structural Similarity Index Metric (SSIM) and Multi-Scale SSIM (MSSSIM) [10].

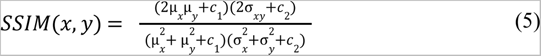

Where:

- μ*_x_* and μ*_y_* are the pixel-wise mean values of x and y
- σ*_xy_* covariance between x and y
- 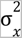 and 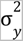 are the variances of x and y
- *c*_1_ = (*k*_1_*L*)^2^ and *c*_2_ = (*k*_2_*L*)^2^, *L* = 2*^#bits_per_pixel^* and *k*_1_ = 0.01 and *k*_2_ = 0.03

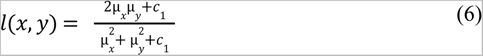

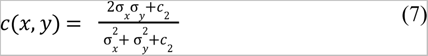

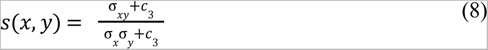

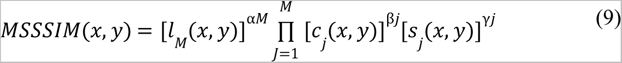

Where:

- *l*(*x*,*y*), *c*(*x*,*y*) and *s*(*x*,*y*) are luminance, contrast and structural similarity measures.
- β_1_ = γ_1_ = 0. 0448, β_2_ = γ_2_ = 0. 2856, β_3_ = γ_3_ = 0. 3001, β_4_ = γ_4_ = 0. 2363, β_5_ = γ_5_ = α_5_ = 0. 1333
- Best results for M = 2

Even though this improves the results, OS is still present, potentially leading to loss of information in high-frequency regions such as cerebellar GM [31]. Learned Perceptual Image Patch Similarity (LPIPS) is a similarity metric that makes use of already trained classification neural networks that were trained with naturalistic images, comparing each of the filters at every depth of the neural network using L2 [33]. Then, they fine-tuned the weights of the network using a human-judgement-based loss so that the metric corresponds better to the human vision behaviour.

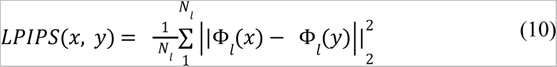

This approach produces a similarity metric that is closer to the human judgment and is more sensitive to blurriness that might perceptually alter the image but produce seemingly high MAE values [33]. The already-trained network can detect features such as edges and textures. As a result, LPIPS can compare each of these features at a deeper level than just comparing two images without being filtered by a network. These characteristics make deep features losses interesting tools to explore for our application of interest. Following these results, we used the first four convolutional layers of a pre-trained network VGG-16 (named after Visual Geometry Group network with 16 layers of weights) as feature extraction, i.e., we generated the feature maps for both denoised and ground truth references using the mentioned pre-trained layers, and then computed the L1 distance between every pair of corresponding feature maps [34]:

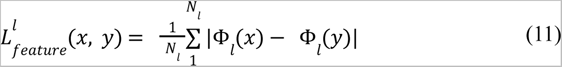

Where *_x_* and *_y_* are the two images to compare, Φ*_l_* are the pre-trained weights of a CNN at a given convolutional layer *l*, and *N_l_* is the number of pixels at *l*. We used the *vgg16* pre trained weights from the *torchvision.models* module, with *l* ∈ {1}.

Finally, the well-known PSNR is a metric that makes pixel-wise comparisons between two images in a logarithmic scale. It can be found in works aiming for image enhancement and more recently in super-resolution tasks. It is computed as follows:

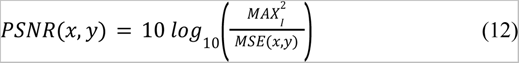

Where *MAX_I_* is the maximum voxel intensity value (e.g., 255 for uint8 image format), and *MSE*(*x*,*y*) is defined in eq. (4).

### 2.7. Data preprocessing

As previously discussed, all MRIs inherently contain a certain amount of noise. This is also the case for all the datasets to be used for training and testing FONDUE (with the exemption of the IXI dataset, which contains virtually zero noise as noted in [32]). The first step in our preprocessing pipeline is to generate surrogates of noise-free MRIs by using a state-of-the-art denoising algorithm. In this case, we used the PRI-NLPCA algorithm proposed by [8], which is a method that effectively filters noisy MRIs to produce denoised images. Using denoised images as ground truth is standard practice and we considered PRI-NLPCA to be a good choice for generating noise-free surrogates since it has been shown to outperform other widely known methods like BM4D and NLM [8]. We did not perform bias field correction at any point since we wanted the network to learn denoising of both, corrected and uncorrected MRIs. The following additional steps were performed:

- We resampled the volumes to isotropic size (equivalent to the minimum voxel size in the 3 dimensions) and number of voxels (equivalent to the maximum dimension in the 3 dimensions) in all 3 dimensions. In other words, an image with voxel size of v1, v2, and v3 and dimensions of H, W, and C was resampled to an image with isotropic voxel size of min(v1,v2,v3) mm^3^ and dimension of max(H, W, C). Resampling operations were made using PyTorch interpolate method using ‘bilinear’ mode, without aligning corners and without recomputing the scaling factor.
- We transformed the volumes to have the standard RAS orientation.
- We robustly rescaled the voxel values to *uchar* values in range [0, 255] using a histogram-based approach with 1000 bins in the histogram, cropping the one thousandth most intense voxels to account for possible outliers. The algorithm was originally developed by Martin Reuter and extracted from [17] under an Apache License, Version 2.0. Finally, we linearly cast the images to type *single* and range [0, 1].

### 2.8. Data augmentation

To introduce invariance to noise levels, we added variable synthetic Rician noise to the training images as input with sigma values ranging randomly between 1.0 and 9.0% (for epochs 1 to 7) and sigma values of 0.0 to 3% for epochs 8 and 9. The variable field consisted of an x3 multiplier in the center of the image volume and ×1 in the outermost voxels, using cubic interpolation for intermediate voxels (i.e. maximum standard deviation was 27%). Furthermore, we performed random horizontal flips and random rotations ranging between [-90, 90] degrees. Lastly, similar to exSA in FastSurferVINN (Henschel et al., 2022), we introduced scale augmentation where instead of bilinearly resampling the images to a fixed resolution inside the network, resolution was modified by adding a µ value to the rescaling value. We set µ as a random number with mean of zero and standard deviation of 0.1. This introduction of scale augmentation can help the network achieve voxel-size invariance.

### 2.9. Training settings

Using a one-cycle learning rate combined with high learning rates and annealing (such as cosine annealing), one could achieve training convergence considerably faster [35]. This approach (called Super Convergence) has since been used to reduce training time in various applications [35], [36]. According to the Super Convergence strategy, OneCycleLR scheduler from PyTorch library was used with the following parameters: max_lr = 0.01 cosine annealing, cycle_momentum = True, pct_start = 0.075, div_factor = 10 and final_div_factor = 100 for the first 5 epochs. For epochs 6 to 9, we used a constant learning rate of 1e-5, using variable low noise levels for the epochs 8 and 9 as described above. AdamW optimizer was used with β_1_ = 0.9 and β_2_ = 0.99. As reported in [37], AdamW is superior to Adam and SGD when adopting a Super Convergence one-cycle learning rate policy. We also explored optimizer settings to ensure optimal performance. PSNR, SSIM, MS-SSIM, and LPIPS metrics were computed for all validation experiments and compared across the four denoising methods. Finally, we used PyTorch’s automatic mixed precision through *torch.cuda.amp.autocast().* This strategy automatically detects the variables and operations that can be done using single-precision instead of double-precision. This strategy enables lighter and faster training while maintaining similar accuracy to using double-precision variables and nearly halves CUDA memory usage.

### 2.10. Data and Code availability statement

Data used for training and validation was obtained from open access repositories, namely ADNI (https://ida.loni.usc.edu/login.jsp?project=ADNI), ABIDE-II (http://fcon_1000.projects.nitrc.org/indi/abide/databases.html), HCP (https://www.humanconnectome.org/study/hcp-young-adult/document/1200-subjects-data-release), IXI (https://brain-development.org/ixi-dataset/), LA5c (https://openneuro.org/datasets/ds000030/versions/00016), MIRIAD (https://www.ucl.ac.uk/drc/research/research-methods/minimal-interval-resonance-imaging-alzheimers-disease-miriad), and UH_NNN (https://openneuro.org/datasets/ds003563/versions/1.0.1).

The source code for FONDUE, installation instructions, and all pre-trained models are available at https://github.com/waadgo/FONDUE.

## 3. Results

We assessed the performance of FONDUE in three different settings: 1. Using held-out images from the same datasets FONDUE was trained on; 2. Using high-quality ground truth images of high resolution (0.5 mm^3^) acquired by averaging 20 repetitions of the same image for two participants; and 3. Using multi-resolution (1 mm^3^, 0.5 mm^3^, and 0.25 mm^3^) data acquired with previously unseen field of strength (7T) and scanner model obtained from the UH_1000, UH_500, and UH_250 respectively [38]. Finally, processing times were compared across different methods to provide a benchmark for performance speed of FONDUE.

### Lambda values

Table 2 presents the obtained *λ_N_* values. The variability across these estimated values indicates that the noise sub-candidates contribute in a different manner to produce the best possible noise final estimation.

**Table 2:**
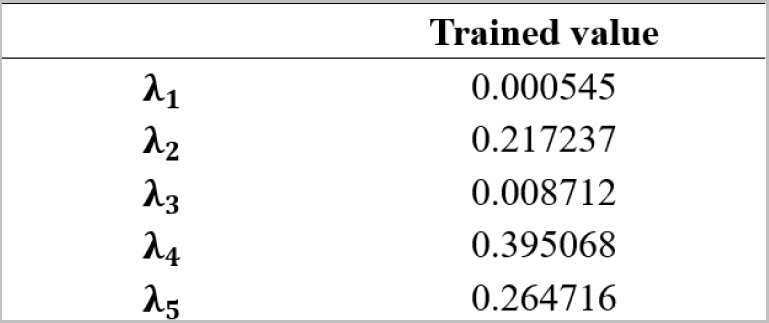
Obtained values for *λ_N_* during training

### 3.1. Generalization on unseen images from datasets used for training

We used 20 images per dataset for ABIDE-II, ADNI1, HCP, IXI, and LA5c and 14 images for MIRIAD dataset to test how well FONDUE generalizes to never-seen-before volumes with similar characteristics (i.e. scanner vendor and acquisition parameters) to those it has been trained on, to ensure the method is not overfitting to the images used for training. On this test, FONDUE obtained the best overall performance for 4 out of the 6 datasets tested (see Tables S.1–S.6 in the supplementary materials). Notably, we can see that PSNR did not always agree with structural similarity and perceptual similarity metrics, which were more likely to fluctuate proportionally.

#### PSNR

FONDUE generally outperformed the compared methods in terms of this metric with the exception of ABIDE-II and ADNI1 datasets, for which ODCT and PRI-NLM were the best two methods and performed similarly to one another. For 9% noise levels, PRI-NLM slightly outperformed FONDUE for HCP and IXI.

#### SSIM and MS-SSIM

Analogously to PSNR, FONDUE overall outperformed the compared methods. The metrics for ABIDE-II and ADNI1 seemed to show a smaller gap between the best-performing methods and FONDUE.

#### LPIPS

For 3 and 7% noise levels, FONDUE achieved the best or second-best performance across all datasets. For 9% of noise level, FONDUE achieved the best performance for LA5c, MIRIAD, and IXI. For very low noise levels, such as 1%, PRI-NLPCA and ODCT achieved the best performance.

Figures 3 and 4 show the performance of all models on representative ABIDE and HCP participants, respectively. FONDUE was able to effectively produce high-quality denoised images even at very high noise levels, even on datasets in which it did not clearly outperform other methods. Furthermore, other methods tended to blur the image structure compared to FONDUE.

**Figure 3:**
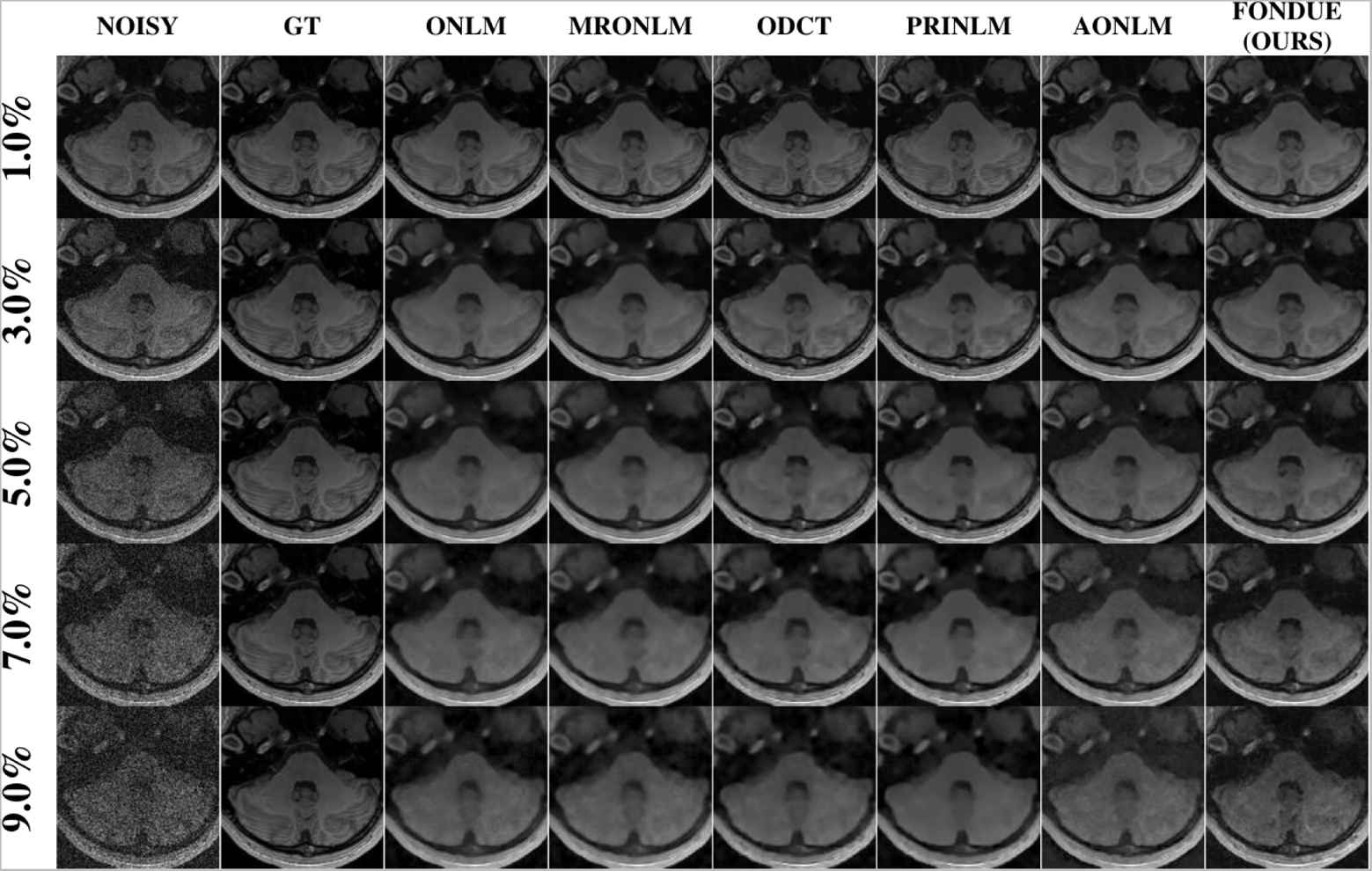
Visual comparison of performance of a generalization subject from dataset ABIDE-II on different Rician noise levels.

**Figure 4.**
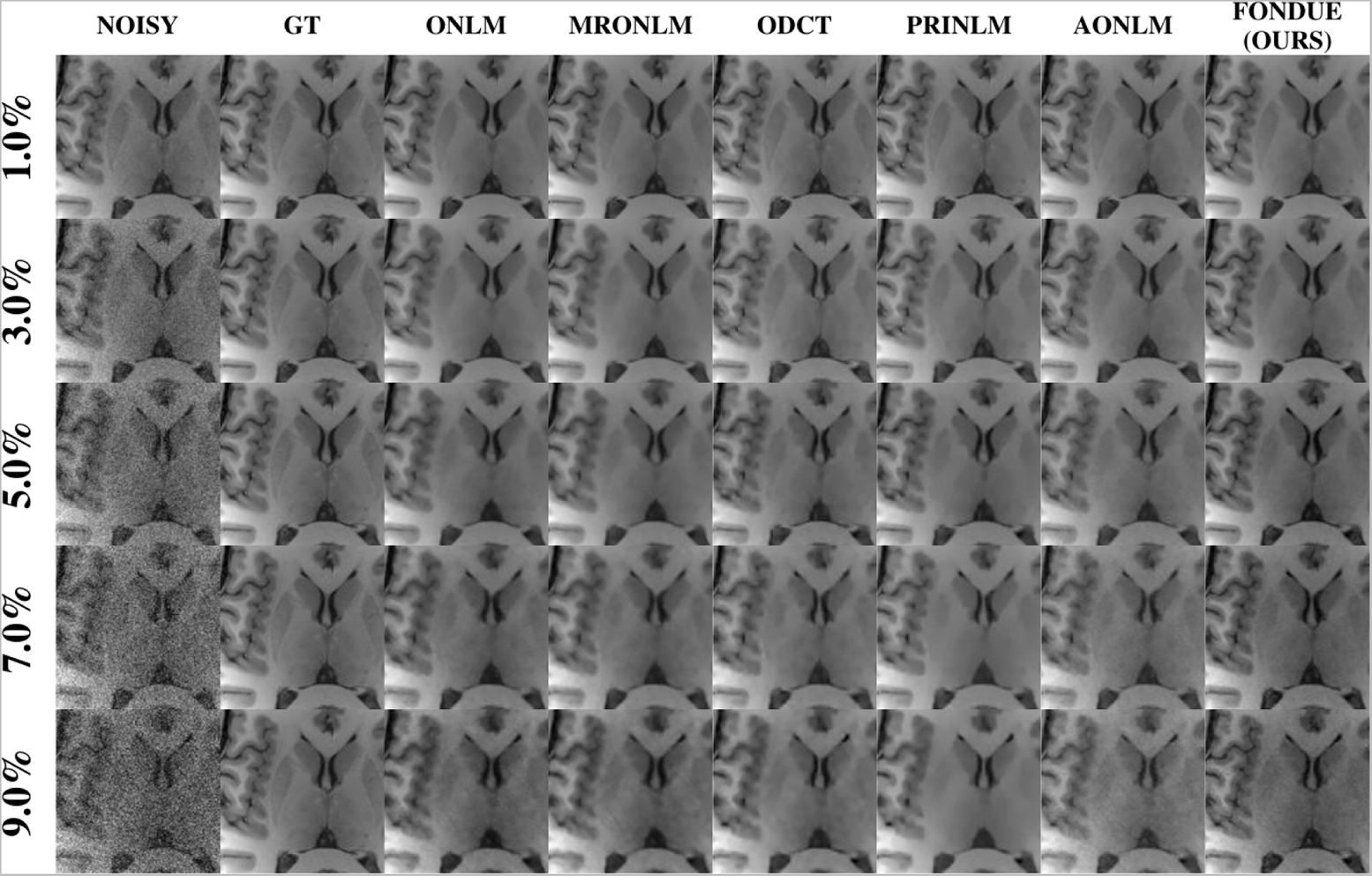
Visual comparison of performance of a generalization subject from dataset HCP on different Rician noise levels.

### 3.2. Generalization on a 0.5 mm^3^ dataset with a high-SNR ground truth

Table 3 summarizes the results of our experiments assessing the performance of FONDUE on a 0.5 mm^3^ dataset with a high-SNR ground truth acquired by averaging 20 repetitions of the same image. Quality metrics indicate an increase in similarity with the reference image as the number of repetitions used for averaging increased. For this test, we used the AONLM-filtered version of the 20-repetition high-SNR image for each of the 2 subjects as reference (ground truth). These results are only provided for AONLM and FONDUE, as the other four techniques (i.e. ONLM, ODCT, MRONLM, and PRINLM) were not able to successfully process these high-resolution images. All four methods use a function called “MRINoiseEstimation”, packed into a mex64 file as a pre-processing stage. The output to this function is parsed as input to other specific secondary functions that produce the denoised results. This function failed with the error *“error during noise estimation”* for high-resolution images. Therefore, we were unable to complete comparisons between FONDUE and ONLM, ODCT, MRONLM, and PRINLM.

**Table 3:** Comparison between denoising single repetition images with AONLM and FONDUE (bottom two rows) vs averaging multiple acquisitions of the same image (rows 1 to 20) on CUSTOM_0.5_20rep dataset. Note: Reference image was the average of 20 repetitions with final filtering using AONLM. Results for subject 1 of the validation set.

| *Image (number of repetitions averaged)* | *LPIPS* | *PSNR (dB)* | *SSIM* | *MSSSIM* |
| --- | --- | --- | --- | --- |
| *1_rep* | 0.057899 | 32.70963 | 0.956913 | 0.988711 |
| *2_rep* | 0.056249 | 33.71358 | 0.960204 | 0.990061 |
| *3_rep* | 0.040658 | 35.76178 | 0.974077 | 0.993824 |
| *4_rep* | 0.033142 | 36.23365 | 0.980061 | 0.995083 |
| *5_rep* | 0.028555 | 36.20465 | 0.98352 | 0.995632 |
| *6_rep* | 0.025292 | 37.04908 | 0.985979 | 0.996347 |
| *7_rep* | 0.024474 | 39.87815 | 0.987925 | 0.99744 |
| *8_rep* | 0.022488 | 39.73384 | 0.989509 | 0.997656 |
| *9_rep* | 0.021088 | 41.11415 | 0.99067 | 0.998034 |
| *10_rep* | 0.019866 | 42.25164 | 0.991503 | 0.998274 |
| *11_rep* | 0.018868 | 43.43498 | 0.992154 | 0.998475 |
| *12_rep* | 0.017914 | 43.39051 | 0.992747 | 0.998522 |
| *13_rep* | 0.017095 | 44.41048 | 0.993083 | 0.998643 |
| *14_rep* | 0.015969 | 45.85579 | 0.993886 | 0.998902 |
| *15_rep* | 0.014906 | 47.01599 | 0.994647 | 0.999088 |
| *16_rep* | 0.014043 | 48.00182 | 0.995686 | 0.999326 |
| *17_rep* | 0.013237 | 48.97971 | 0.995686 | 0.999326 |
| *18_rep* | 0.012577 | 49.61897 | 0.996093 | 0.9994 |
| *19_rep* | 0.012066 | 50.20783 | 0.996341 | 0.999453 |
| *20_rep* | 0.011628 | 50.57085 | 0.996609 | 0.999494 |
| *FONDUE (mean ± stdev)* | 0.015 ± 0.0016 | 39.65 ± 1.73 | 0.991 ± 0.0013 | 0.997 ± 0.0006 |
| *AONLM (mean ± stdev)* | 0.024 ± 0.0028 | 39.45 ± 2.14 | 0.986 ± 0.0018 | 0.996 ± 0.0008 |

FONDUE showed a very good performance in denoising both images (see Figure 5 for an example). For subject 1, FONDUE showed similarity with the reference comparable to averaging ~15 co-registered repetitions of the image in terms of LPIPS, ~7 in terms of PSNR, ~9 in terms of SSIM, and ~6 in MSSSIM. For comparison, AONLM performed similarly to averaging ~10, ~6, ~6, and ~5 in terms of LPIPS, PSNR, SSIM, and MSSIM, respectively. For subject 2, FONDUE showed a similarity to the reference image comparable with averaging ~20, ~16, ~13, and ~10 repetitions in terms of LPIPS, PSNR, SSIM, and MSSSIM. AONLM performed similarly to averaging ~8, ~9, ~7, and ~6 repetitions, respectively. In addition to consistently outperforming AONLM, FONDUE showed less variability reflected in the standard deviations reported in Tables 3 and S.7 (last two rows). Interestingly, PSNR was the only metric not monotonically improving with the number of repetitions averaged (e.g., from 4 to 5, from 7 to 8, and from 11 to 12 repetitions included in the average, PSNR decreased), whereas LPIPS, SSIM, and MSSSIM always showed an improvement.

**Figure 5:**
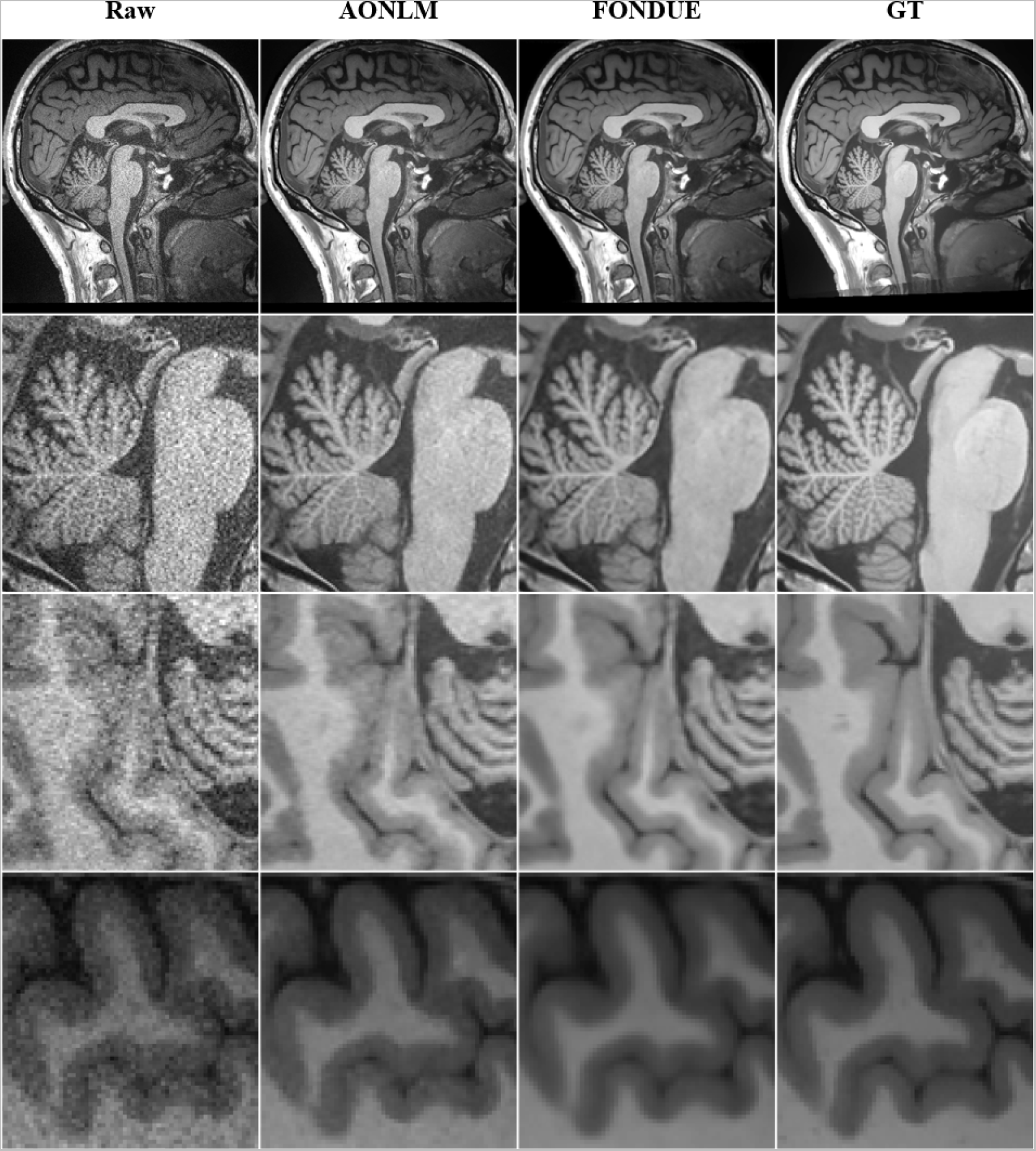
Visual comparison of FONDUE and AONLM on CUSTOM_0.5_20rep dataset

### 3.3. Qualitative results on UH_250, UH_500, and UH_1000

For this validation, we tested the denoising capabilities of FONDUE using the same subject’s brain images acquired at three different resolutions to assess how well FONDUE generalizes to unseen resolutions and magnetic field strengths (i.e. 7T). While noise levels were very low for low resolution images (1.0mm^3^ isotropic), as expected, they incremented with the increase in image resolution. At 0.25 mm^3^, there is a considerable amount of noise that makes it difficult to distinguish the true anatomy on the image. FONDUE was able to effectively filter-out the natural noise from the images at every resolution, while preserving important high-frequency information (Figure 5). Both AONLM and FONDUE considerably enhanced the 0.25 mm^3^ image, making it easier to visualize the structures captured by the scanner. However, based on visual assessments, FONDUE produced a higher quality denoised image, with more consistent textures on both low and high-frequency regions, in particular noticeable in GM color uniformity and in the contrast between cerebellar GM and WM. Similar to the 0.5 mm^3^ data, ONLM, ODCT, MRONLM, and PRINLM were not able to successfully process the 0.25 mm^3^ data, and we were thus unable to include them in these comparisons.

**Figure 6.**
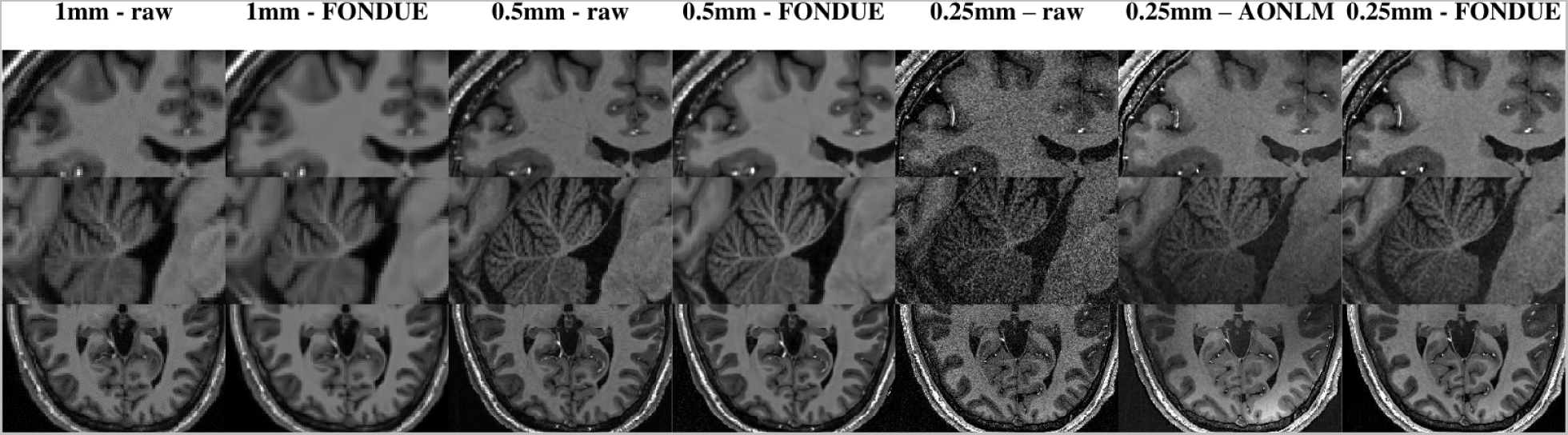
UH_NNN dataset denoised at three different resolutions by FONDUE and comparison vs AONLM at 0.25mm isotropic voxel size. FONDUE can denoise at unseen resolutions and unseen field strength (7T).

### 3.4. Processing times

Table 4 shows average processing times for AONLM and FONDUE for 0.5 mm^3^ isotropic voxel size images using an image of size 448×448×448 voxels. FONDUE was slightly faster (~12%) when used on a CPU, and significantly faster (~30 times) when using a GPU.

**Table 4.**
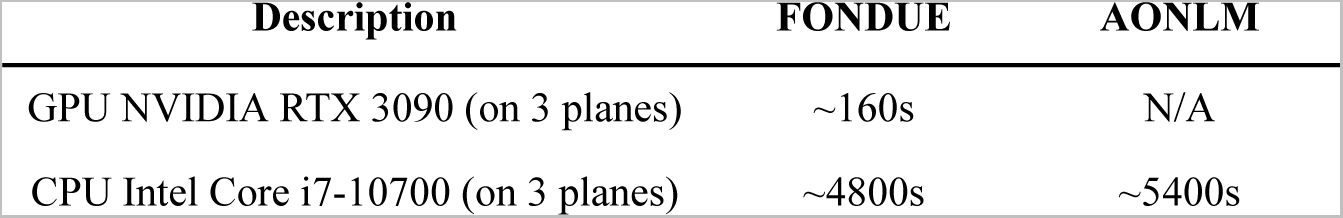
Processing times of FONDUE & AONLM on the used system.

## 4. Discussion

In this paper, we have proposed FONDUE, a human brain structural MRI denoising method capable of handling multiple resolutions, scanner models, and field strengths, even outside the resolutions for which it was trained. The proposed method outperformed or matched the state-of-the-art methods when compared to reference surrogates (silver standard). The usage of the proposed perceptual loss based on VGG-16 resulted in a good balance between denoising and detail preservation.

The training time was ~6h per epoch using a batch size of 2, turning the proposed method into an affordable one. Total training time was less than 3 days on the-still domestic grade-NVIDIA RTX 3090 thanks to the super convergence strategy. Including 0% noise levels in training was crucial for optimal results, since the network learned how to not degrade noise-free images, which is a potential issue when not considering training using noise-free images. Similarly, the inclusion of automatic mixed precision (AMP) during the training made it feasible to denoise images with varying resolutions-including ultra-high resolution images from the UH_250 dataset-using an entry-level NVIDIA RTX 3060 GPU, since AMP uses around half the GPU memory and reduces the computational cost.

We further report on the unreliability of using solely PSNR as a quality metric, as it did not monotonically increase as more repetitions were considered for averaging on the CUSTOM_0.5 dataset (Tables 3 and S.7). More complex metrics such as LPIPS, SSIM, and MS-SSIM monotonically improved as more repetitions were included for the high-SNR average image. Therefore, our observations agree with the recent literature on the need for using similarity metrics that can be more similar to human visual perception.

As opposed to the traditional approaches that use UNet/UNet++-like architectures [11], [21] in which the filters number increases while going deeper into the encoder part of the network, our approach managed to keep low to very low GPU memory usage by keeping the same number of filters throughout the network. Furthermore, our experiments showed that not all the sub-candidates “YN” produced by the sub-networks should have the same weight when averaging to produce the final result (see Table 2).

In our experiments, FONDUE did not perform optimally for images from ABIDE-II and ADNI1. It is worth noting that these datasets both had non-isotropic voxel sizes, whereas the rest of datasets contained only MR images acquired with isotropic voxel sizes. This difference in performance levels was likely caused by the interpolation step in the pre-processing stage as well as the trained weights being optimized for images acquired using isotropic voxel sizes. However, the results using non-isotropic images were still of high quality and comparable to the other reported methods. Furthermore, recent studies considering structural MRIs mostly use isotropic resolutions in the scanner acquisition parameters. Our validations further showed that FONDUE generalized well on healthy and diseased cohorts of varied demographics.

Overall, FONDUE was able to provide high-quality denoised images across a range of settings in significantly shorter processing times than state-of-the-art denoising methods when using a GPU. Future work will include training using other loss functions such as FLIP and E-LPIPS, training specific weights for each anatomical plane instead of using the axial weights for the three anatomical planes, validating FONDUE on other structural modalities of brain MRI such as T2w, T2*, FLAIR and PD, and training a 2D-only version of FONDUE, to optimize it for usage with non-isotropic images.

## Acknowledgments

Dr. Dadar reports receiving research funding from the Healthy Brains for Healthy Lives (HBHL), Alzheimer Society Research Program (ASRP), Douglas Research Centre (DRC), and Natural Sciences and Engineering Research Council of Canada (NSERC).

## Supplementary Materials

**Figure S.1.**
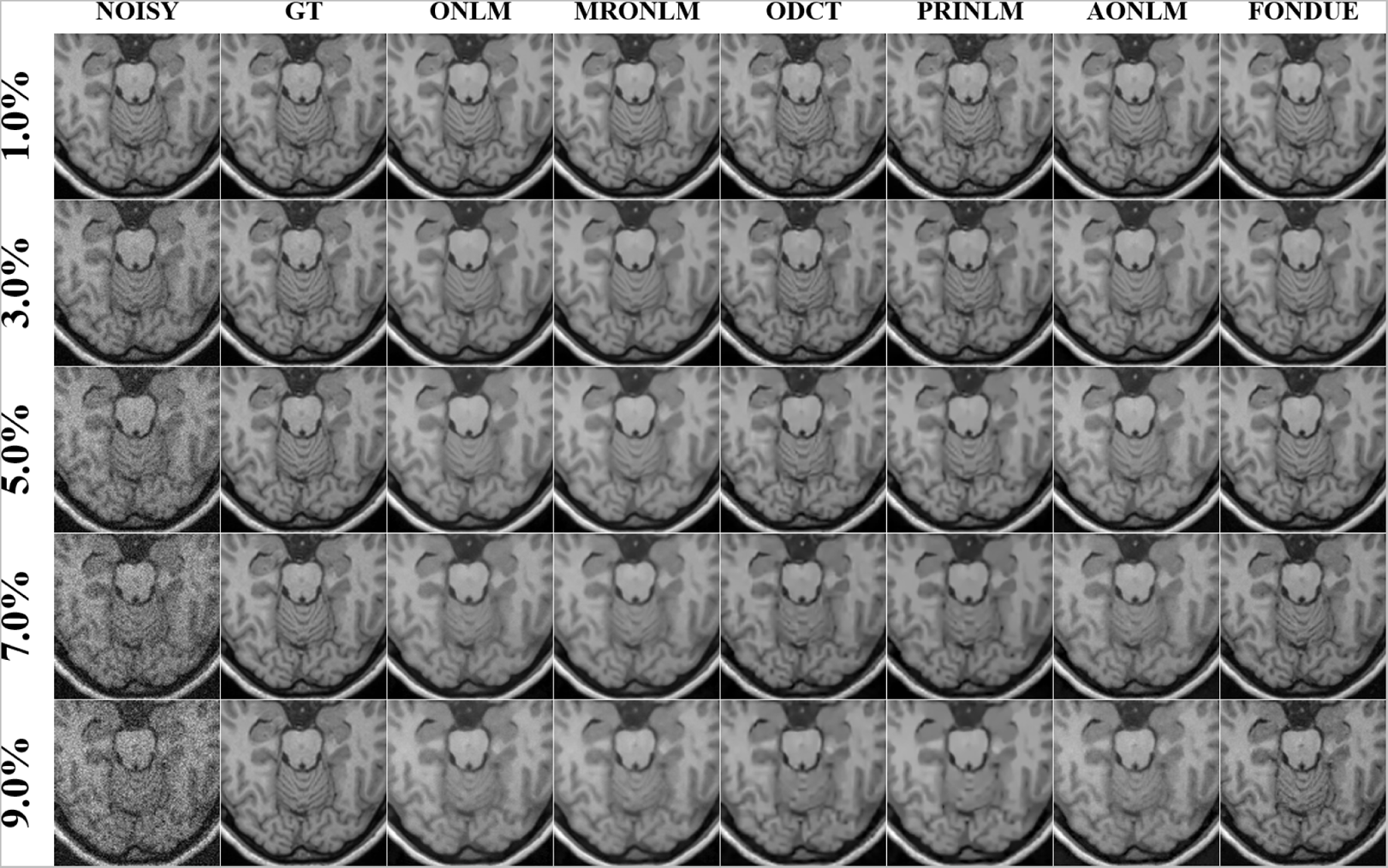
Visual comparison of the performance of a generalization subject from dataset IXI on different Rician noise levels

**Figure S.2.**
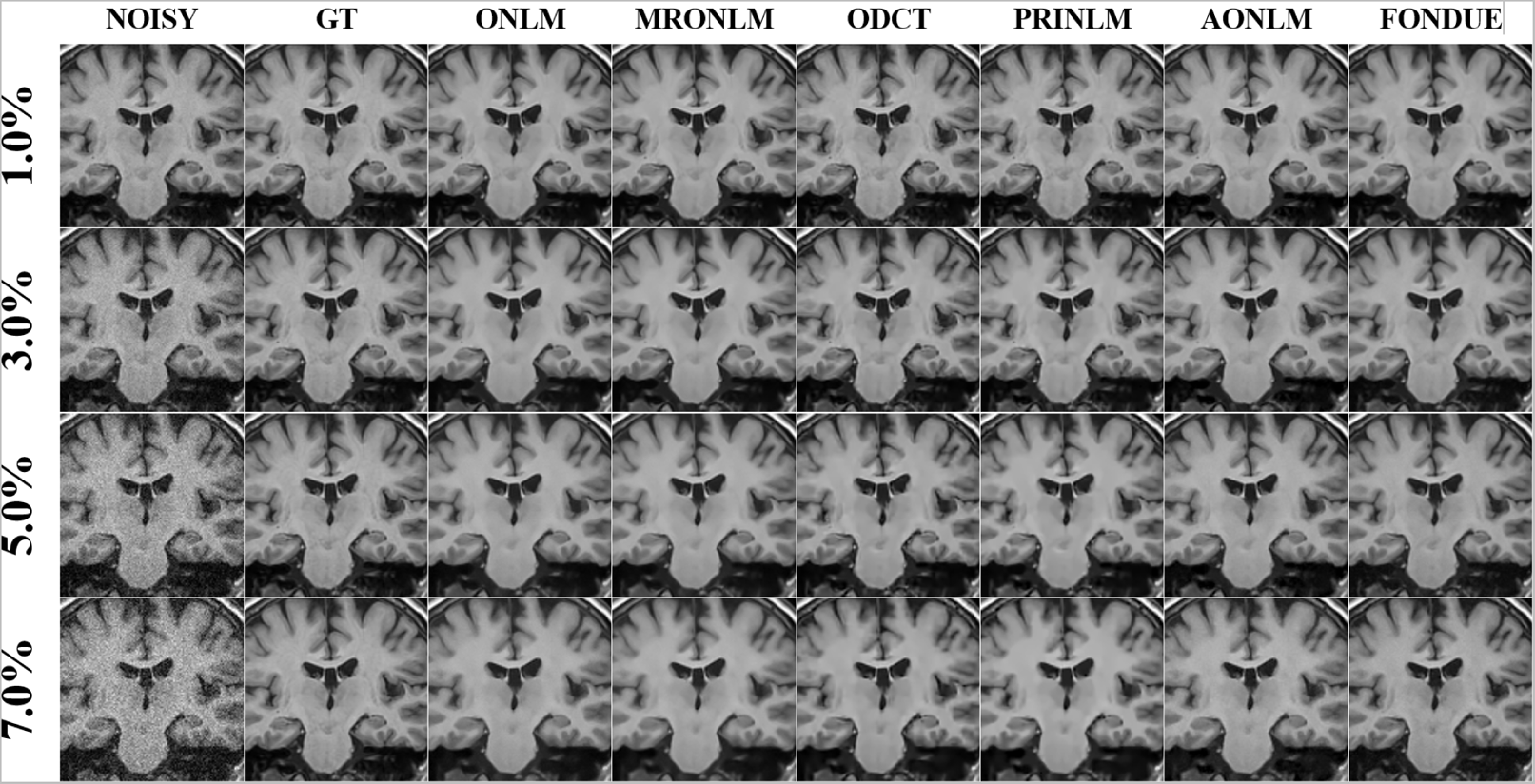
Visual comparison of the performance of a generalization subject from dataset MIRIAD on different Rician noise levels.

**Figure S.3.**
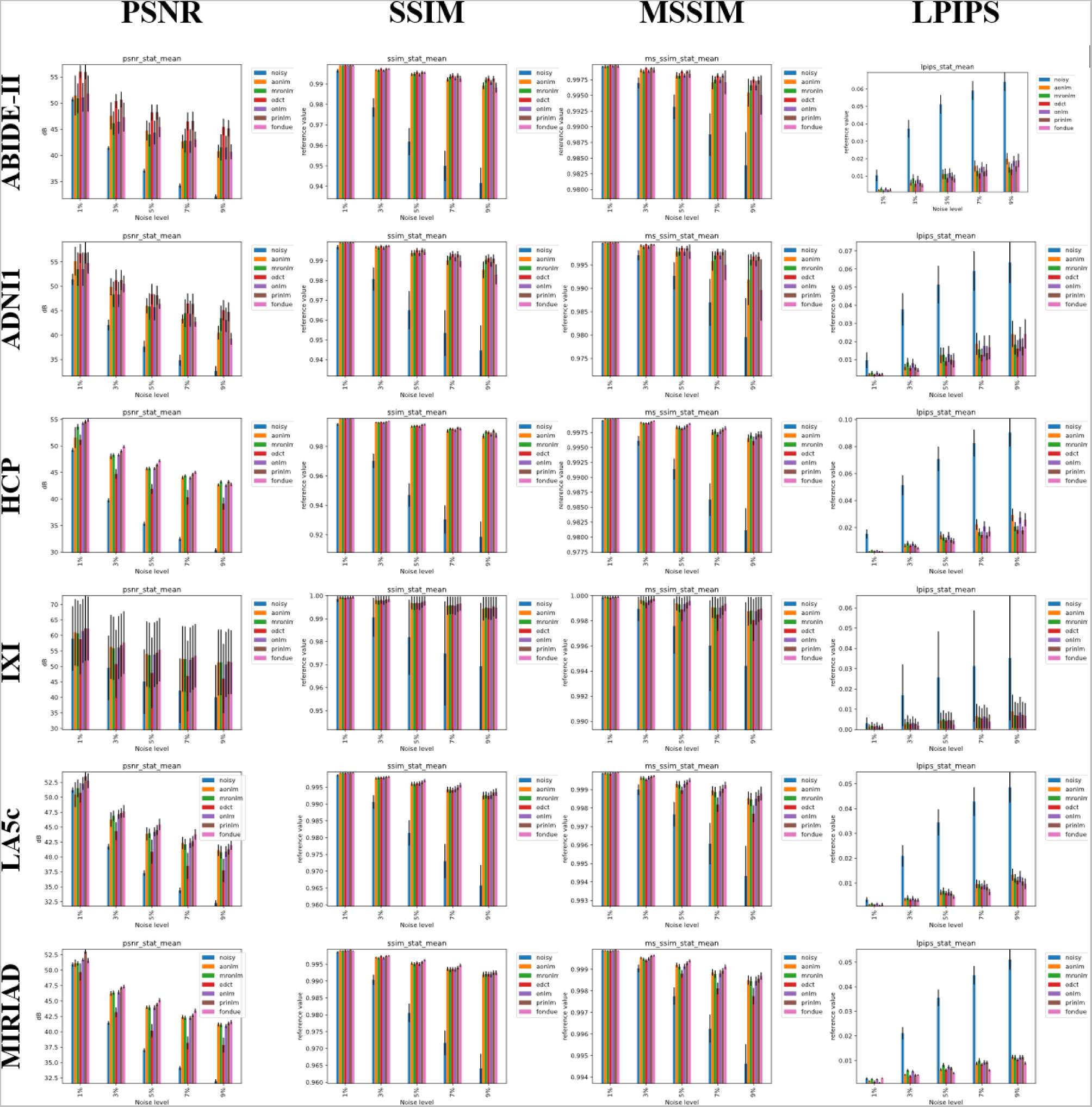
Visual comparison of the performance of a generalization subject from dataset MIRIAD on different Rician noise levels.

**Table S.1:** ABIDE-II Generalization set results

|  |  | 1% | 3% | 5% | 7% | 9% | Average |
| --- | --- | --- | --- | --- | --- | --- | --- |
| PSNR | Noisy | 50.7 | 41.4 | 37.0 | 34.2 | 32.1 | 39.1 |
|  | AONLM | 51.5 | 47.5 | 44.7 | 42.6 | 40.8 | 45.4 |
|  | MRONLM | 50.9 | 46.2 | 44.0 | 42.8 | 41.5 | 45.1 |
|  | ODCT | **56.0** | 50.4 | 48.2 | 46.5 | 45.4 | **49.3** |
|  | ONLM | 51.1 | 46.5 | 44.4 | 42.7 | 41.5 | 45.2 |
|  | PRINLM | 56.0 | **50.7** | **48.3** | **46.6** | **45.2** | **49.3** |
|  | FONDUE | 51.8 | 47.3 | 45.4 | 43.1 | 40.7 | 45.7 |
| SSIM | Noisy | 0.99641 | 0.97850 | 0.96188 | 0.95000 | 0.94156 | 0.96567 |
|  | AONLM | 0.99890 | 0.99685 | 0.99466 | 0.99218 | 0.98911 | 0.99434 |
|  | MRONLM | 0.99892 | 0.99654 | 0.99485 | 0.99352 | 0.99196 | 0.99516 |
|  | ODCT | **0.99917** | 0.99729 | **0.99565** | **0.99415** | **0.99274** | **0.99580** |
|  | ONLM | 0.99894 | 0.99655 | 0.99457 | 0.99280 | 0.99071 | 0.99471 |
|  | PRINLM | 0.99916 | 0.99730 | 0.99562 | 0.99413 | 0.99260 | 0.99576 |
|  | FONDUE | 0.99893 | **0.99732** | 0.99544 | 0.99250 | 0.98817 | 0.99447 |
| MS-SSIM | Noisy | 0.999583 | 0.996975 | 0.993172 | 0.988769 | 0.983854 | 0.99247 |
|  | AONLM | 0.999593 | 0.998981 | 0.998189 | 0.997076 | 0.995442 | 0.997856 |
|  | MRONLM | 0.999623 | 0.998783 | 0.998108 | 0.997488 | 0.996594 | 0.998119 |
|  | ODCT | **0.999843** | **0.999349** | **0.9988** | **0.998169** | **0.997503** | **0.998733** |
|  | ONLM | 0.999636 | 0.998837 | 0.998156 | 0.997443 | 0.99657 | 0.998128 |
|  | PRINLM | 0.999841 | **0.999349** | 0.998769 | 0.998158 | 0.997397 | 0.998703 |
|  | FONDUE | 0.999632 | 0.999035 | 0.998422 | 0.997084 | 0.995055 | 0.997845 |
| LPIPS | Noisy | 0.0105 | 0.0372 | 0.0512 | 0.0591 | 0.0641 | 0.0444 |
|  | AONLM | 0.0020 | 0.0062 | 0.0111 | 0.0158 | 0.0200 | 0.0110 |
|  | MRONLM | 0.0029 | 0.0086 | 0.0112 | 0.0130 | 0.0150 | 0.0101 |
|  | ODCT | **0.0016** | 0.0057 | 0.0089 | **0.0115** | **0.0139** | **0.0083** |
|  | ONLM | 0.0027 | 0.0079 | 0.0117 | 0.0152 | 0.0186 | 0.0112 |
|  | PRINLM | 0.0017 | 0.0061 | 0.0098 | 0.0126 | 0.0157 | 0.0092 |
|  | FONDUE | 0.0023 | **0.0048** | **0.0086** | 0.0134 | 0.0190 | 0.0096 |

**Table S.2:** ADNI1 Generalization set results

|  |  | 1% | 3% | 5% | 7% | 9% | Average |
| --- | --- | --- | --- | --- | --- | --- | --- |
| PSNR | Noisy | 51.4 | 42.1 | 37.7 | 34.9 | 32.6 | 39.7 |
|  | AONLM | 55.2 | 49.9 | 46.0 | 43.3 | 40.5 | 47.0 |
|  | MRONLM | 53.5 | 48.4 | 45.8 | 44.5 | 43.4 | 47.1 |
|  | ODCT | **56.7** | 51.2 | **48.4** | **46.5** | **45.1** | **49.6** |
|  | ONLM | 53.5 | 48.3 | 45.7 | 44.4 | 43.1 | 47.0 |
|  | PRINLM | **56.9** | **51.3** | **48.2** | **46.4** | **44.8** | **49.5** |
|  | FONDUE | 54.8 | 50.5 | 46.4 | 42.7 | 39.2 | 46.7 |
| SSIM | Noisy | 0.996837 | 0.980748 | 0.96504 | 0.953471 | 0.944612 | 0.968142 |
|  | AONLM | 0.999019 | 0.996868 | 0.993777 | 0.990182 | 0.985388 | 0.993047 |
|  | MRONLM | 0.998966 | 0.996395 | 0.994016 | 0.9922 | 0.9904 | 0.994396 |
|  | ODCT | **0.99917** | 0.997352 | **0.995342** | **0.993278** | **0.991224** | **0.995273** |
|  | ONLM | 0.998964 | 0.996383 | 0.993802 | 0.991676 | 0.989386 | 0.994042 |
|  | PRINLM | **0.999163** | 0.997355 | 0.995298 | 0.993176 | 0.991019 | 0.995202 |
|  | FONDUE | 0.999044 | **0.997415** | 0.994387 | 0.989744 | 0.983071 | 0.992732 |
| MS-SSIM | Noisy | 0.999627 | 0.997087 | 0.992663 | 0.986976 | 0.979578 | 0.991186 |
|  | AONLM | 0.999782 | 0.999144 | 0.997809 | 0.995827 | 0.99183 | 0.996879 |
|  | MRONLM | 0.999667 | 0.998804 | 0.997809 | 0.997016 | 0.996088 | 0.997877 |
|  | ODCT | **0.999837** | **0.999352** | **0.998639** | **0.997772** | **0.996808** | **0.998482** |
|  | ONLM | 0.999668 | 0.998806 | 0.997773 | 0.997008 | 0.995955 | 0.997842 |
|  | PRINLM | **0.999841** | **0.999346** | 0.998569 | **0.99776** | 0.996763 | 0.998456 |
|  | FONDUE | 0.99975 | 0.999295 | 0.997821 | 0.994984 | 0.989657 | 0.996302 |
| LPIPS | Noisy | 0.009663 | 0.037768 | 0.051471 | 0.058938 | 0.063643 | 0.044297 |
|  | AONLM | 0.001988 | 0.005935 | 0.012427 | 0.018731 | 0.024187 | 0.012653 |
|  | MRONLM | 0.002764 | 0.008448 | 0.012743 | 0.015671 | 0.018296 | 0.011584 |
|  | ODCT | **0.001673** | 0.005334 | **0.009015** | **0.012603** | **0.016148** | **0.008955** |
|  | ONLM | 0.002693 | 0.008025 | 0.013129 | 0.017525 | 0.021415 | 0.012557 |
|  | PRINLM | 0.001779 | 0.005758 | 0.009963 | 0.013655 | 0.017108 | 0.009653 |
|  | FONDUE | 0.002051 | **0.004325** | **0.009567** | 0.01683 | 0.024217 | 0.011398 |

**Table S.3.** HCP Generalization set results

|  |  | 1% | 3% | 5% | 7% | 9% | Average |
| --- | --- | --- | --- | --- | --- | --- | --- |
| PSNR | Noisy | 49.2 | 39.7 | 35.3 | 32.5 | 30.3 | 37.4 |
|  | AONLM | 51.6 | 48.0 | 45.7 | 44.0 | 42.7 | 46.4 |
|  | MRONLM | 53.6 | 48.3 | 45.8 | 44.4 | 43.2 | 47.0 |
|  | ODCT | 51.1 | 44.7 | 41.9 | 40.3 | 39.1 | 43.4 |
|  | ONLM | 54.2 | 48.3 | 45.7 | 44.0 | 42.5 | 46.9 |
|  | PRINLM | 54.6 | 49.0 | 46.4 | 44.7 | **43.3** | 47.6 |
|  | FONDUE | **54.9** | **49.8** | **47.2** | **45.0** | 42.8 | **47.9** |
| SSIM | Noisy | 0.995067 | 0.970242 | 0.947081 | 0.9304 | 0.918549 | 0.952268 |
|  | AONLM | 0.998763 | 0.99631 | 0.993545 | 0.990582 | 0.987137 | 0.993267 |
|  | MRONLM | 0.998833 | 0.996139 | 0.993836 | 0.991942 | 0.989948 | 0.99414 |
|  | ODCT | 0.998765 | 0.996506 | 0.994095 | 0.991815 | 0.989606 | 0.994157 |
|  | ONLM | 0.998849 | 0.996074 | 0.993492 | 0.990986 | 0.988154 | 0.993511 |
|  | PRINLM | 0.998931 | 0.996794 | 0.994702 | **0.992711** | **0.990659** | **0.994759** |
|  | FONDUE | **0.998937** | **0.997126** | **0.994982** | 0.991982 | 0.987586 | 0.994123 |
| MS-SSIM | Noisy | 0.999471 | 0.996115 | 0.991382 | 0.986283 | 0.981157 | 0.990882 |
|  | AONLM | 0.999749 | 0.999128 | 0.998365 | 0.997542 | 0.996548 | 0.998266 |
|  | MRONLM | 0.99976 | 0.999008 | 0.998317 | 0.997705 | 0.996991 | 0.998356 |
|  | ODCT | 0.999747 | 0.999027 | 0.998116 | 0.997145 | 0.996096 | 0.998026 |
|  | ONLM | 0.999774 | 0.999025 | 0.99833 | 0.997657 | 0.996856 | 0.998328 |
|  | PRINLM | 0.999813 | 0.999277 | 0.998645 | 0.997967 | **0.997184** | 0.998577 |
|  | FONDUE | **0.999818** | **0.99942** | **0.998935** | **0.998242** | 0.997137 | **0.99871** |
| LPIPS | Noisy | 0.015336 | 0.051481 | 0.070863 | 0.082619 | 0.090393 | 0.062138 |
|  | AONLM | 0.002271 | 0.006786 | 0.014256 | 0.022291 | 0.029395 | 0.015 |
|  | MRONLM | 0.003047 | 0.008411 | 0.012754 | 0.016603 | 0.020808 | 0.012325 |
|  | ODCT | 0.002382 | 0.006544 | 0.010823 | 0.01466 | 0.018303 | 0.010542 |
|  | ONLM | 0.002968 | 0.007979 | 0.013927 | 0.020908 | 0.027423 | 0.014641 |
|  | PRINLM | 0.00222 | 0.006619 | 0.010729 | **0.014217** | **0.017921** | **0.010341** |
|  | FONDUE | **0.002155** | **0.004767** | **0.009942** | 0.017113 | 0.025837 | 0.011963 |

**Table S.4:** IXI Generalization set results

|  |  | **1%** | **3%** | **5%** | **7%** | **9%** | **Average** |
| --- | --- | --- | --- | --- | --- | --- | --- |
| PSNR | Noisy | 59.0 | 49.5 | 45.1 | 42.2 | 40.0 | 47.2 |
|  | AONLM | 61.0 | 56.3 | 54.0 | 52.4 | 51.2 | 55.0 |
|  | MRONLM | 60.7 | 55.8 | 53.7 | 52.4 | 51.3 | 54.8 |
|  | ODCT | 58.8 | 50.8 | 47.9 | 46.9 | 46.1 | 50.1 |
|  | ONLM | 61.4 | 56.1 | 53.8 | 52.1 | 50.6 | 54.8 |
|  | PRINLM | **62.3** | 56.9 | 54.4 | 52.8 | **51.6** | 55.6 |
|  | FONDUE | 62.2 | **57.7** | **55.4** | **53.4** | 51.4 | **56.0** |
| SSIM | Noisy | 0.99857 | 0.99062 | 0.981942 | 0.97484 | 0.969305 | 0.983055 |
|  | AONLM | 0.999288 | 0.998113 | 0.996905 | 0.995657 | 0.994357 | 0.996864 |
|  | MRONLM | 0.999262 | 0.997954 | 0.996773 | 0.99581 | 0.994947 | 0.996949 |
|  | ODCT | 0.99914 | 0.998147 | 0.996913 | 0.995801 | 0.994672 | 0.996935 |
|  | ONLM | 0.999281 | 0.997974 | 0.996757 | 0.995632 | 0.994503 | 0.996829 |
|  | PRINLM | 0.999361 | 0.998295 | 0.997289 | 0.99632 | **0.995305** | 0.997314 |
|  | FONDUE | **0.99937** | **0.998574** | **0.997741** | **0.996611** | 0.994764 | **0.997412** |
| MS-SSIM | Noisy | 0.999862 | 0.998953 | 0.997587 | 0.996037 | 0.994413 | 0.99737 |
|  | AONLM | 0.999902 | 0.999665 | 0.999385 | 0.999084 | 0.998763 | 0.99936 |
|  | MRONLM | 0.999886 | 0.999605 | 0.99932 | 0.999066 | 0.998814 | 0.999338 |
|  | ODCT | 0.999849 | 0.999461 | 0.998915 | 0.9985 | 0.998073 | 0.99896 |
|  | ONLM | 0.999894 | 0.999621 | 0.99934 | 0.999066 | 0.998776 | 0.99934 |
|  | PRINLM | **0.999915** | 0.999708 | 0.999469 | 0.99921 | 0.998912 | 0.999443 |
|  | FONDUE | **0.999915** | **0.999765** | **0.999594** | **0.999364** | **0.998986** | **0.999525** |
| LPIPS | Noisy | 0.003078 | 0.016961 | 0.025673 | 0.031295 | 0.035262 | 0.022454 |
|  | AONLM | 0.001347 | 0.00269 | 0.004373 | 0.006623 | 0.008988 | 0.004804 |
|  | MRONLM | 0.001896 | 0.003633 | 0.004962 | 0.006056 | 0.007146 | 0.004739 |
|  | ODCT | 0.001441 | 0.002748 | 0.004223 | 0.005531 | 0.006854 | 0.004159 |
|  | ONLM | 0.001834 | 0.003358 | 0.004677 | 0.006425 | 0.008426 | 0.004944 |
|  | PRINLM | **0.00111** | 0.002877 | 0.004386 | 0.005707 | 0.007159 | 0.004248 |
|  | FONDUE | 0.001442 | **0.001934** | **0.002478** | **0.003859** | **0.006795** | **0.003301** |

**Table S.5:** LA5c Generalization set results

|  |  | 1% | 3% | 5% | 7% | 9% | Average |
| --- | --- | --- | --- | --- | --- | --- | --- |
| PSNR | Noisy | 59.0 | 49.5 | 45.1 | 42.2 | 40.0 | 47.2 |
|  | AONLM | 61.0 | 56.3 | 54.0 | 52.4 | 51.2 | 55.0 |
|  | MRONLM | 60.7 | 55.8 | 53.7 | 52.4 | 51.3 | 54.8 |
|  | ODCT | 58.8 | 50.8 | 47.9 | 46.9 | 46.1 | 50.1 |
|  | ONLM | 61.4 | 56.1 | 53.8 | 52.1 | 50.6 | 54.8 |
|  | PRINLM | **62.3** | 56.9 | 54.4 | 52.8 | **51.6** | 55.6 |
|  | FONDUE | 62.2 | **57.7** | **55.4** | **53.4** | 51.4 | **56.0** |
| SSIM | Noisy | 0.998578 | 0.990616 | 0.981398 | 0.973021 | 0.965735 | 0.98187 |
|  | AONLM | 0.999189 | 0.997668 | 0.996032 | 0.99432 | 0.992535 | 0.995949 |
|  | MRONLM | 0.999248 | 0.997774 | 0.995985 | 0.994226 | 0.992686 | 0.995984 |
|  | ODCT | 0.999195 | 0.997876 | 0.996096 | 0.994131 | 0.992413 | 0.995942 |
|  | ONLM | 0.999272 | 0.997826 | 0.996139 | 0.994404 | 0.992688 | 0.996066 |
|  | PRINLM | **0.999369** | 0.998045 | 0.996522 | 0.994942 | 0.993411 | 0.996458 |
|  | FONDUE | 0.999307 | **0.998106** | **0.996992** | **0.995645** | **0.99355** | **0.99672** |
| MS-SSIM | Noisy | 0.999868 | 0.998993 | 0.997656 | 0.996073 | 0.994326 | 0.997383 |
|  | AONLM | 0.999872 | 0.999619 | 0.999294 | 0.998939 | 0.998538 | 0.999252 |
|  | MRONLM | 0.999888 | 0.999621 | 0.999263 | 0.998865 | 0.998466 | 0.999221 |
|  | ODCT | 0.999851 | 0.999537 | 0.998965 | 0.998186 | 0.997692 | 0.998846 |
|  | ONLM | 0.9999 | 0.999638 | 0.999295 | 0.998914 | 0.998501 | 0.99925 |
|  | PRINLM | **0.999921** | 0.999697 | 0.999395 | 0.999045 | 0.998662 | 0.999344 |
|  | FONDUE | **0.99992** | **0.999728** | **0.999513** | **0.999239** | **0.998785** | **0.999437** |
| LPIPS | Noisy | 0.003288 | 0.020995 | 0.034484 | 0.042853 | 0.04858 | 0.03004 |
|  | AONLM | 0.001354 | 0.003515 | 0.006286 | 0.009671 | 0.013405 | 0.006846 |
|  | MRONLM | 0.001714 | 0.004063 | 0.006863 | 0.009536 | 0.011947 | 0.006825 |
|  | ODCT | 0.001196 | 0.003242 | 0.005975 | 0.008677 | 0.011016 | 0.006021 |
|  | ONLM | 0.001697 | 0.003871 | 0.006314 | 0.009303 | 0.012602 | 0.006757 |
|  | PRINLM | **0.001024** | 0.0032 | 0.005828 | 0.008346 | 0.01061 | 0.005802 |
|  | FONDUE | 0.001421 | **0.00314** | **0.004547** | **0.0064** | **0.009723** | **0.005046** |

**Table S.6:** MIRIAD Validation set results

|  |  | 1% | 3% | 5% | 7% | 9% | Average |
| --- | --- | --- | --- | --- | --- | --- | --- |
| PSNR | Noisy | 50.9 | 41.4 | 37.0 | 34.1 | 32.0 | 39.1 |
|  | AONLM | 51.1 | 46.2 | 44.0 | 42.4 | 41.2 | 45.0 |
|  | MRONLM | 51.1 | 46.3 | 43.9 | 42.3 | 41.1 | 45.0 |
|  | ODCT | 49.7 | 43.2 | 40.2 | 38.2 | 37.8 | 41.8 |
|  | ONLM | 51.7 | 46.4 | 43.9 | 42.2 | 40.9 | 45.0 |
|  | PRINLM | **53.0** | 47.1 | 44.4 | 42.7 | 41.4 | 45.7 |
|  | FONDUE | 51.6 | **47.3** | **45.1** | **43.4** | **41.6** | **45.8** |
| SSIM | Noisy | 0.998599 | 0.990395 | 0.980556 | 0.971657 | 0.964066 | 0.981055 |
|  | AONLM | 0.999007 | 0.996978 | 0.995254 | 0.993605 | 0.991953 | 0.995359 |
|  | MRONLM | 0.998996 | 0.996755 | 0.994987 | 0.99339 | 0.992085 | 0.995243 |
|  | ODCT | 0.999 | 0.997366 | 0.995424 | 0.993562 | 0.992037 | 0.995478 |
|  | ONLM | 0.999009 | 0.996794 | 0.995009 | 0.993444 | 0.991947 | 0.995241 |
|  | PRINLM | **0.999252** | 0.997396 | 0.995605 | 0.993986 | **0.992509** | 0.995749 |
|  | FONDUE | 0.998883 | **0.997516** | **0.99617** | **0.994673** | 0.992451 | **0.995939** |
| MS-SSIM | Noisy | 0.999875 | 0.999041 | 0.997755 | 0.99625 | 0.994624 | 0.997509 |
|  | AONLM | 0.999869 | 0.999536 | 0.999202 | 0.998858 | 0.998503 | 0.999194 |
|  | MRONLM | 0.999859 | 0.999474 | 0.999124 | 0.998779 | 0.998454 | 0.999138 |
|  | ODCT | 0.999844 | 0.999403 | 0.9988 | 0.998114 | 0.997759 | 0.998784 |
|  | ONLM | 0.999869 | 0.999488 | 0.999138 | 0.998805 | 0.998461 | 0.999152 |
|  | PRINLM | **0.999913** | 0.999619 | 0.999278 | 0.998927 | 0.998563 | 0.99926 |
|  | FONDUE | 0.999863 | **0.999641** | **0.999395** | **0.999113** | **0.99869** | **0.999341** |
| LPIPS | Noisy | 0.002642 | 0.021147 | 0.03553 | 0.044699 | 0.051079 | 0.03102 |
|  | AONLM | 0.001605 | 0.004259 | 0.006359 | 0.00878 | 0.011489 | 0.006498 |
|  | MRONLM | 0.002472 | 0.00598 | 0.008081 | 0.009874 | 0.011241 | 0.00753 |
|  | ODCT | 0.001297 | 0.003597 | 0.006108 | 0.008369 | 0.010306 | 0.005935 |
|  | ONLM | 0.002376 | 0.005598 | 0.00736 | 0.009193 | 0.011315 | 0.007168 |
|  | PRINLM | **0.000975** | **0.004037** | 0.006829 | 0.009187 | 0.011339 | 0.006473 |
|  | FONDUE | 0.002756 | 0.004064 | **0.00491** | **0.006038** | **0.008849** | **0.005323** |

**Table S.7:** Comparison between denoising a single repetition with AONLM and FONDUE (two bottom rows) vs averaging multiple acquisitions of the same image (row 1 to row 20): CUSTOM_0.5 dataset. Note: The reference image used was the average of 20 repetitions with a final filtering using AONLM. Results for subject 2 of the validation set.

| *Image (number of repetitions averaged)* | *LPIPS* | *PSNR (dB)* | *SSIM* | *MSSSIM* |
| --- | --- | --- | --- | --- |
| *1_rep* | 0.0548 | 23.78113 | 0.948708 | 0.974982 |
| *2_rep* | 0.040766 | 26.30681 | 0.964412 | 0.985924 |
| *3_rep* | 0.033366 | 27.79532 | 0.971802 | 0.989935 |
| *4_rep* | 0.0286 | 29.39115 | 0.976705 | 0.992717 |
| *5_rep* | 0.025199 | 29.69427 | 0.979702 | 0.993445 |
| *6_rep* | 0.022809 | 29.9983 | 0.981929 | 0.994043 |
| *7_rep* | 0.021007 | 30.76593 | 0.984026 | 0.994969 |
| *8_rep* | 0.019102 | 31.59893 | 0.985916 | 0.995777 |
| *9_rep* | 0.017769 | 32.60824 | 0.987448 | 0.996504 |
| *10_rep* | 0.0165 | 34.31008 | 0.988752 | 0.997322 |
| *11_rep* | 0.015535 | 34.76009 | 0.989697 | 0.997568 |
| *12_rep* | 0.014627 | 35.89668 | 0.990567 | 0.997953 |
| *13_rep* | 0.013877 | 36.70565 | 0.991297 | 0.998191 |
| *14_rep* | 0.01307 | 35.7144 | 0.991969 | 0.9981 |
| *15_rep* | 0.012215 | 35.38514 | 0.992649 | 0.998121 |
| *16_rep* | 0.011601 | 37.35103 | 0.993386 | 0.998599 |
| *17_rep* | 0.01101 | 37.45049 | 0.993913 | 0.998683 |
| *18_rep* | 0.010476 | 36.58092 | 0.994309 | 0.998593 |
| *19_rep* | 0.009991 | 38.00277 | 0.994803 | 0.998875 |
| *20_rep* | 0.009514 | 38.08047 | 0.995157 | 0.99893 |
| *FONDUE (mean ± stdev)* | 0.0086 ± 0.001 | 37.33 ± 2.05 | 0.991 ± 0.0008 | 0.997 ± 0.0003 |
| *AONLM (mean ± stdev)* | 0.019 ± 0.003 | 32.14 ± 3.14 | 0.983 ± 0.002 | 0.994 ± 0.0009 |

